# GEOlimma: Differential Expression Analysis and Feature Selection Using Pre-Existing Microarray Data

**DOI:** 10.1101/693564

**Authors:** Liangqun Lu, Kevin A. Townsend, Bernie J. Daigle

## Abstract

**Background:** Differential expression and feature selection analyses are essential steps for the development of accurate diagnostic/prognostic classifiers of complicated human diseases using transcriptomics data. These steps are particularly challenging due to the curse of dimensionality and the presence of technical and biological noise. A promising strategy for overcoming these challenges is the incorporation of pre-existing transcriptomics data in the identification of differentially expressed (DE) genes. This approach has the potential to improve the quality of selected genes, increase classification performance, and enhance biological interpretability. While a number of methods have been developed that use pre-existing data for differential expression analysis, existing methods do not leverage the identities of experimental conditions to create a robust metric for identifying DE genes.

**Results:** In this study, we propose a novel differential expression and feature selection method—GEOlimma—which combines pre-existing microarray data from the Gene Expression Omnibus (GEO) with the widely-applied Limma method for differential expression analysis. We first quantify differential gene expression across 2481 pairwise comparisons from 602 curated GEO Datasets, and we convert differential expression frequencies to DE prior probabilities. Genes with high DE prior probabilities show enrichment in cell growth and death, signal transduction, and cancer-related biological pathways, while genes with low prior probabilities were enriched in sensory system pathways. We then applied GEOlimma to four differential expression comparisons within two human disease datasets and performed differential expression, feature selection, and supervised classification analyses. Our results suggest that use of GEOlimma provides greater experimental power to detect DE genes compared to Limma, due to its increased effective sample size. Furthermore, in a supervised classification analysis using GEOlimma as a feature selection method, we observed similar or better classification performance than Limma given small, noisy subsets of an asthma dataset.

**Conclusions:** Our results demonstrate that GEOlimma is a more effective method for differential gene expression and feature selection analyses compared to the standard Limma method. Due to its focus on gene-level differential expression, GEOlimma also has the potential to be applied to other high-throughput biological datasets.

## Background

DNA microarrays and RNA sequencing (RNA-Seq) have become indispensable experimental tools for characterizing the effects of biological interventions on genome-wide gene expression (“transcriptomics”) [1] [2]. Applications of these tools have been transformative in many areas of biological research, including cancer biology, biomarker discovery, and drug target identification [3] [4] [5]. These applications often involve differential expression analysis: the isolation of differentially expressed (DE) genes between healthy and disease conditions. Knowledge of DE genes facilitates the discovery of causative genes and gene pathways for a disease of interest. For example, many studies of carcinogenesis focus on identifying the genes directly responsible for promoting cancer occurrence (“driver genes”) out of all DE genes [6]. Furthermore, DE gene identification is an important first step for disease biomarker discovery. The discovery of biomarkers from transcriptomics data typically involves selecting the most discriminative genes between a healthy and diseased state or between different disease states [7]. A comprehensive list of DE genes provides a biologically plausible set of candidates for these discriminative genes and can greatly streamline the search [8]. Common applications of transcriptomics-derived biomarkers include predicting diagnosis, prognosis, and therapeutic response for a disease of interest through a process known as supervised classification [9]. In this context, DE gene identification can be viewed as a means of performing feature selection for classification. In general, feature selection is a process for dimensionality reduction that removes redundant or irrelevant features (genes), reduces classification model complexity, and improves classification performance [10].

Despite their widespread use for DE gene identification, transcriptomics data are notorious for their inclusion of technical and biological noise [11]. This noise complicates differential expression analysis by reducing the accuracy of DE gene identification relative to other assays (e.g., real-time or quantitative PCR [12]), lowering the reproducibility of experiments conducted on different platforms [13], and reducing the statistical power associated with the detection of DE genes at a particular fold change [14]. A straightforward strategy for mitigating the effects of noise is to increase the number of replicates assayed (“sample size”) for each condition of interest. However, this practice can be cost prohibitive or even impossible for conditions with limited sample availability. Furthermore, even with larger sample sizes, transcriptomics data pose a considerable challenge to feature selection methods due to the curse of dimensionality. Specifically, it is well known that optimal fitting of classification models (including the selection of features) breaks down when the feature dimensionality is substantially larger than the sample size [15].

One promising solution for the above challenges is to incorporate prior biological knowledge into differential expression and feature selection analyses [16]. This Bayesian approach can mitigate problems associated with a small sample size [17], while also improving biological interpretability of the resulting DE genes/features [10]. Prior biological knowledge for transcriptomics data can take several forms, including pre-existing transcriptomics data from other studies, data from complementary high-throughput assays (e.g., chromatin immunoprecipitation or protein-protein interactions), and gene functional annotation (e.g., Gene Ontology [18] [19] or KEGG [20] [21]). For the purposes of this study, we will focus on the first type of knowledge, although we note that analytical methods are available to incorporate the other types as well [22] [23]. Thanks to functional genomics repositories like the Gene Expression Omnibus (GEO)[24] [25] and ArrayExpress [26], transcriptomics data from over 2.5 million samples are publicly available. Furthermore, the size of this resource is growing exponentially, with numbers of samples in GEO doubling every 3-4 years.

Over the last 15 years, a number of methods have been developed that use prior knowledge in the form of transcriptomics data to inform differential expression analyses [27] [28] [29] [30] [31]. However, these methods typically either ignore the identities of the many experimental conditions in the pre-existing data, or they do not leverage these identities to create a rigorous statistical metric for identifying DE genes. For example, the SVD Augmented Gene expression Analysis Tool (SAGAT) uses singular value decomposition (SVD) to extract transcriptional modules from pre-existing DNA microarray data [27]. These modules, which contain no information regarding assayed conditions, are then incorporated into a statistical analog of the two-sample t-test to improve the accuracy of DE gene identification. In contrast, a very recent study made direct use of the experimental conditions in pre-existing data to characterize empirical prior probabilities of differential expression [31]. However, although these prior probabilities were predictive of differential expression patterns, they were not explicitly utilized in a Bayesian statistical framework for identifying DE genes. Relatedly, although there have been many studies contributing novel or adapted feature selection methodologies for classification of biomedical data [32] [33] [34] [35] [36], to our knowledge no method combines an experimental condition-aware analysis of pre-existing data with a statistically principled means of feature selection.

To address these shortcomings, we propose a novel differential expression and feature selection approach—GEOlimma—that leverages pre-existing GEO-derived transcriptomics data. As described below, our proposed method modifies the popular Linear Models for Microarray and RNA-Seq Data (“Limma”) method [37][38]. Specifically, GEOlimma incorporates empirical prior probabilities of differential expression (DE prior probabilities) in a Bayesian statistical test for DE genes. We first describe the computation and biological characterization of DE prior probabilities from a large collection of pre-existing DNA microarray experiments from GEO. Next, we apply GEOlimma and Limma to four benchmark differential expression comparisons from two validation datasets. Our results demonstrate a substantial increase in experimental power for identifying DE genes due to use of GEOlimma. Finally, we explore GEOlimma’s ability to improve feature selection for classification across the four benchmark comparisons.

## Methods

### GEOlimma Method Formulation

We developed the GEOlimma method by combining the widely-used differential expression (DE) analysis method Limma, which is typically used to analyze gene expression microarray and RNA-seq data and assess differential expression between biological conditions. Limma uses empirical Bayesian methods to provide stable DE predictions, which is particularly useful when the number of sample replicates is small. However, one simplifying assumption made by Limma is that the DE prior probabilities for each gene are identical (set 0.01 by default). GEOlimma combines the Bayesian nature of Limma with gene-level DE prior probabilities calculated from large-scale microarray datasets to better select genes that are biologically relevant to a comparison of interest.

The Gene Expression Omnibus (GEO) is a public data repository for high-throughput gene expression data including microarray and RNA-seq data [25]. GEO DataSets (GDS) are a subset of the repository that store curated gene expression datasets, along with the original data (GEO Series) and experimental platform information. GPL570, also known as the `HG-U133_Plus_2` Affymetrix Human Genome U133 Plus 2.0 Array, is one of the best-represented human genome microarray platforms in GEO, with 149,049 samples available (as of June 7, 2019). GPL570 measures over 47,000 human transcripts, which consist of the Human Genome U133 Set plus 6,500 additional genes. In this study, we downloaded all 602 GPL570 GEO DataSets (GDS) (current as of June 7, 2019). Specifically, for each dataset we obtained normalized, log-transformed expression values at the probeset level. We then mapped these probesets to the non-redundant Entrez Gene IDs (provided by the Bioconductor R package hgu133plus2.db) and obtained gene-level expression values by computing medians across any probe sets mapping to the same gene. With the minimum requirement of 5 samples in each group, we performed pairwise DE analysis among the largest possible collection of non-overlapping sample groups from each GDS experiment. Specifically, for each DE comparison, we applied the Limma moderated t-test [39] (using the “lmFit” and “eBayes” functions) to calculate differential expression p-values for each gene. Given a list of p-values for a particular comparison, we adjusted for multiple hypothesis testing using the Benjamini-Hochberg (BH) procedure [40]. Genes with adjusted p-values (false discovery rates or FDRs) ≤0.05 for a given pairwise comparison were considered DE for that comparison. We calculated the DE frequencies across all comparisons for each gene and converted these frequencies to DE prior probabilities (P(DE)) as follows:

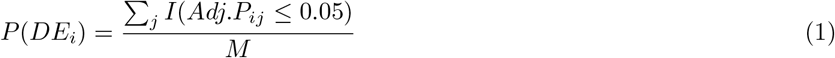

where *i* ∈ {1, …, *N*} indexes each gene, *j* ∈ {1, … , *M*} indexes each comparison, *Adj.P*_*ij*_ represents the FDR for the i-th gene in the j-th comparison, and I(·) is the indicator function.

We chose human asthma and cancer validation datasets present as GEO Series (GSE) but not as GEO DataSets (GDS), in order to avoid double counting data. The asthma dataset [41] consists of 404 total samples (transformed lymphoblastoid cell lines) taken from 268 children afflicted with asthma and 136 healthy children. The cancer dataset [42] consists of 870 total bone marrow samples, of which 202, 164, and 69 are from individuals with acute myeloid leukemia (AML), myelodysplastic syndrome (MDS), and neither AML nor MDS, respectively. We considered the three possible comparisons between these three groups. In total, we evaluated four comparisons: Asthma vs Non-asthma, Nonleukemia vs AML, Nonleukemia vs MDS, and AML vs MDS.

For a given comparison, we compute GEOlimma DE posterior probabilities using Bayes’ theorem:

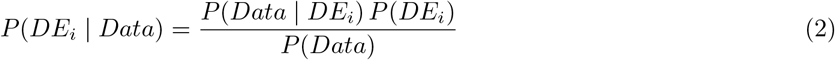

where Data represents the samples making up the given comparison, *P*(*Data* | *DE*_*i*_) denotes the likelihood of the Data, as calculated by limma [37], *P*(*DE*_*i*_) is the previously calculated DE prior probability, and *P*(*Data*) is a normalization constant [37]. Given these posterior probabilities, we then calculate B scores (log odds of DE) for each gene as follows:

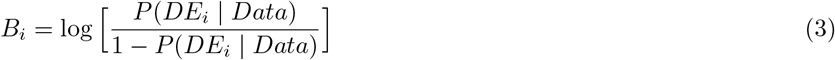

We implemented GEOlimma as modified R functions based on code from the Limma package.

### Enrichment Analysis for Gene Sets

To explore the DE prior probabilities biologically, we conducted KEGG Enrichment Analysis using the R package ClusterProfiler [43]. Specifically, we identified enriched KEGG pathways using the hypergeometric test in both the top and bottom 500 most/least frequently DE genes, separately. Pathways with BH-adjusted p-values less than 0.05 were considered significantly enriched. We used the Pathview R package [44] to visualize the location of DE genes in particular KEGG pathways.

### Differential Expression Analysis

#### Evaluation datasets

As described above, we downloaded the GSEs for two evaluation datasets from GEO. As with the GDS data, we mapped normalized, log-transformed expression values at the probeset level to non-redundant Entrez Gene IDs and consolidated expression values by computing medians across probe sets mapping to the same gene. We included all genes with unique probe mappings (20283 total) for subsequent analyses. For each of the four evaluation comparisons, we performed DE analysis on all samples using both GEOlimma and Limma. Genes were considered DE if their BH-adjusted p-value ≤0.05 (Limma) or their B score exceeded the smallest Limma B score for genes with adjusted p-value ≤0.05 (GEOlimma).

#### Sample Visualization

To visualize samples, we first used Principal Component Analysis (PCA) to reduce the dimensionality of genes as features. We visualized the first two components of PCA. We further applied the t-Distributed Stochastic Neighbor Embedding (t-SNE) method to visualize the first 10 PCA components in 2 dimensions. t-SNE can reduce the dimensionality of data based on conditional probabilities that preserve local similarity. We used a t-SNE implementation that makes Barnes-Hut approximations, allowing it to be applied on large real-world datasets [45]. We set the perplexity to 15, and sample points were colored using the group information.

#### Experimental power

To quantify the performance improvement achieved by GEOlimma vs Limma, we performed DE analysis on small sample size subsets for each comparison. As detailed below, we started with the minimum subset size at which the group proportions for a given comparison could be maintained and generated all non-overlapping sample subsets of this size. We then increased this subset size by the smallest possible sample increment and repeated the generation of subsets. For each sample subset, we first applied both GEOlimma and Limma and ranked genes by their corresponding B scores. Next, using the Limma DE genes previously identified from all samples as the ground truth (see Results section for specific numbers), we applied the R package ROCR [46] to calculate Area under the ROC curves (AUCs) for the B score-ranked genes of each subset. We calculated the performance improvement of GEOlimma over Limma for each subset as the difference in AUC between the two methods. In addition, we converted these AUC improvements into gains in effective sample size by constructing and interpolating from a “standard curve” of mean Limma AUC values calculated across the full range of possible sample sizes. As an example, if GEOlimma delivered an AUC improvement of 0.1 over Limma for a subset of size 10, the GEOlimma effective sample size is simply the sample size of the standard curve corresponding to an AUC value 0.1 higher than the mean Limma AUC value for 10-sample subsets.

### Supervised Classification

We performed supervised classification for each comparison in the evaluation datasets using both GEOlimma and Limma as feature selection methods. Scikit-learn (sklearn) [47] is a Python module implementing machine learning algorithms. It enables various tasks such as dimensionality reduction, classification, regression and model selection. The sklearn classification pipeline involves sequentially applying feature selection, classification, parameter optimization and model selection to yield final classification results. We first used the Python rpy2 module to build a connection between sklearn and the R language, followed by creating customized feature selection methods for Limma and GEOlimma which we compiled into the sklearn pipeline function. For classification training, we first sampled 10 subsets of 40 samples (20 from each of the two groups) at random and selected the 1000 genes with largest variance across these samples. Next, we fed data from each subset to the sklearn pipeline function and performed either Limma or GEOlimma-based feature selection by selecting subsets of 100-1000 genes (in increments of 100) with the highest B scores. We selected the Logistic Regression [48] classifier for classification. We also included L1 and L2 penalties as hyperparameters and applied 10-fold cross validation to train the model and optimize the hyperparameters. We used classification AUC as the criterion to evaluate classification performance. A high AUC represents both high recall and high precision, which translate to low false positive and false negative rates. For classification testing, we sampled an additional 40 samples to evaluate the training models. We used a Wilcoxon signed-rank test to identify significant AUC differences between performing feature selection using Limma or GEOlimma.

## Results

In this study, we developed a gene expression feature selection method, GEOlimma, in which gene-level differential expression (DE) prior probabilities were derived from large-scale microarray data freely available from the Gene Expression Omnibus (GEO). We first explored enriched biological pathways in genes with either high or low DE prior probabilities. We then applied GEOlimma to DE analysis and supervised classification tasks on a collection of four validation datasets.

### Biological Analysis of DE Prior Probabilities

The goal of differential expression analysis is to identify differences in gene expression across biological conditions in order to discover functional genes and pathways involved in a biological process of interest. The Limma method [39] is an empirical Bayesian approach for identifying DE genes that has been widely applied. However, an important limitation of this method is that the prior probabilities for differential expression are set to be constant for all genes. This implies that all genes have the same chance of being expressed differently, which is not biologically realistic [31]. Therefore, we developed and applied GEOlimma, which uses a large collection of GEO datasets to compute gene level DE prior probabilities (see Methods section). We first downloaded the 602 GEO DataSets (GDS) currently available from the GPL570 platform (Affymetrix Human Genome U133 Plus 2.0 Array), followed by performing pairwise DE analysis among the largest possible collection of non-overlapping sample groups (number of samples 5) from each GDS experiment. We identified DE genes using a Benjamini-Hochberg false discovery rate (FDR) threshold of 0.05. By repeating this procedure for every GDS, we calculated DE frequencies for 21025 distinct Entrez genes (20283 genes with unique gene mappings) across all experiments (2481 pairwise comparisons total) and converted these to prior probabilities of DE. Given gene-level DE prior probabilities, we can then compute posterior probabilities of DE for a given biological experiment using Bayes’ theorem. Figure 1 shows the distribution of DE prior probabilities, which ranged between 0.0048 and 0.1769 and appeared to have two modes. The median probability is 0.069, which we note is roughly seven times higher than the default constant prior probability used by Limma (0.01). Figure SFigure1 lists the top most frequently DE genes, including TUBA1A (tubulin alpha 1a), CD24, and SERPINB1 (serpin family B member 1), with DE prior probabilities of 0.1769, 0.1761, and 0.1693, respectively. The three least frequently DE genes were LOC102725116, TMCO5A (transmembrane and coiled-coil domains 5A), and LINC01492 (long intergenic non-protein coding RNA 1492), with DE prior probabilities of 0.0048, 0.0056, and 0.0060, respectively. Generally speaking, we hypothesize that genes with high prior probabilities of DE are more likely to be implicated in human disease and thus could function as biomarkers, while those with low DE prior probabilities represent constitutively expressed genes that are required for the maintenance of basic cellular functions (i.e., housekeeping genes).

**Figure 1.**
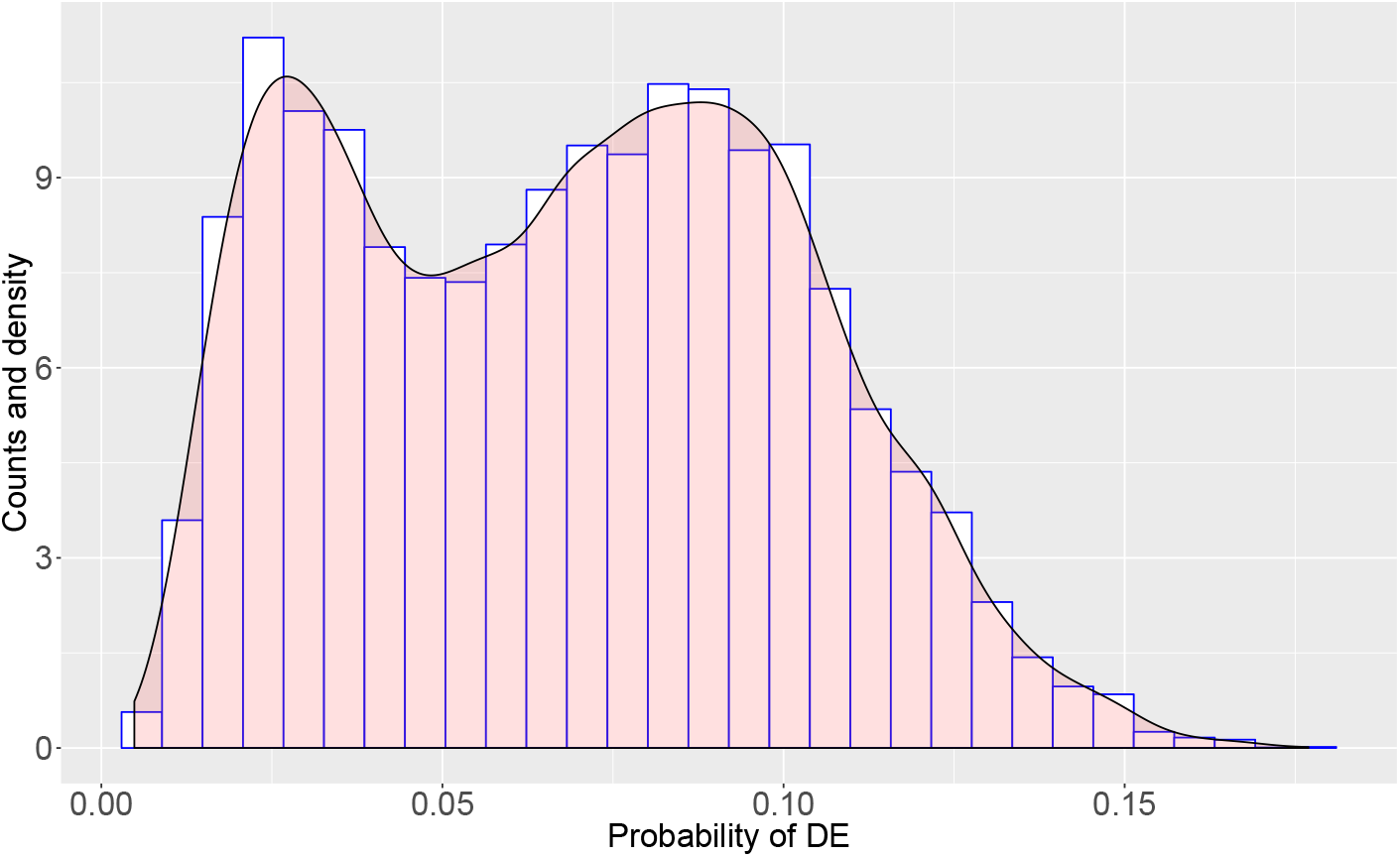
Distribution of DE prior probabilities for 20283 genes, calculated from 2481 pairwise comparisons made within 602 curated GEO Datasets.

In order to improve our biological understanding of the calculated DE prior probabilities, we performed gene set enrichment analysis (GSEA) based on KEGG pathways with the top 500 most and least frequently DE genes, respectively. Table 1 lists significantly enriched pathways (BH-adjusted p-value 0.05), which include 19 pathways from the most frequently DE genes and 4 from the least frequently DE genes. The most significant pathway in the former category is hsa04110: Cell cycle (adjusted p = 7.83E-08); Figure 2 illustrates the frequently DE genes mapped in this pathway. Two additional pathways in this category directly related to cell growth and death include hsa04115: p53 signaling pathway and hsa04210: Apoptosis. We also identified six cancer-specific frequently DE pathways: hsa05222: Small cell lung cancer, hsa05206: MicroRNAs in cancer, hsa05218: Melanoma, hsa05202: Transcriptional misregulation in cancer, hsa05205: Proteoglycans in cancer, and hsa05220: Chronic myeloid leukemia. Finally, the two frequently DE pathways hsa04068: FoxO signaling pathway and hsa04668: TNF signaling pathway function in Signal transduction. We note that signal transduction pathways are involved in cell death mechanisms that function in colorectal carcinogenesis progression [49].

**Table 1.** KEGG Enrichment Analysis of top 500 genes with high and low DE prior probabilities.

| Pathway IDs | Description | GeneRatio | BgRatio | Pvalue | P value adjustment | Count | Source |
| --- | --- | --- | --- | --- | --- | --- | --- |
| hsa04110 | Cell cycle | 21/242 | 124/7528 | 2.94E-10 | 7.83E-08 | 21 | HighPrior |
| hsa05222 | Small cell lung cancer | 13/242 | 93/7528 | 7.47E-06 | 9.94E-04 | 13 | HighPrior |
| hsa04115 | p53 signaling pathway | 11/242 | 72/7528 | 1.61E-05 | 1.43E-03 | 11 | HighPrior |
| hsa05169 | Epstein-Barr virus infection | 18/242 | 201/7528 | 7.63E-05 | 3.57E-03 | 18 | HighPrior |
| hsa05206 | MicroRNAs in cancer | 23/242 | 299/7528 | 8.53E-05 | 3.57E-03 | 23 | HighPrior |
| hsa05218 | Melanoma | 10/242 | 72/7528 | 9.09E-05 | 3.57E-03 | 10 | HighPrior |
| hsa05202 | Transcriptional misregulation in cancer | 17/242 | 186/7528 | 9.40E-05 | 3.57E-03 | 17 | HighPrior |
| hsa04210 | Apoptosis | 14/242 | 136/7528 | 1.12E-04 | 3.71E-03 | 14 | HighPrior |
| hsa05205 | Proteoglycans in cancer | 17/242 | 201/7528 | 2.42E-04 | 7.15E-03 | 17 | HighPrior |
| hsa04068 | FoxO signaling pathway | 13/242 | 132/7528 | 3.05E-04 | 8.11E-03 | 13 | HighPrior |
| hsa05418 | Fluid shear stress and atherosclerosis | 13/242 | 139/7528 | 5.05E-04 | 1.22E-02 | 13 | HighPrior |
| hsa05220 | Chronic myeloid leukemia | 9/242 | 76/7528 | 6.87E-04 | 1.52E-02 | 9 | HighPrior |
| hsa03030 | DNA replication | 6/242 | 36/7528 | 8.98E-04 | 1.84E-02 | 6 | HighPrior |
| hsa05130 | Pathogenic Escherichia coli infection | 7/242 | 55/7528 | 1.77E-03 | 3.36E-02 | 7 | HighPrior |
| hsa04540 | Gap junction | 9/242 | 88/7528 | 1.97E-03 | 3.50E-02 | 9 | HighPrior |
| hsa01524 | Platinum drug resistance | 8/242 | 73/7528 | 2.25E-03 | 3.74E-02 | 8 | HighPrior |
| hsa05167 | Kaposi sarcoma-associated herpesvirus infection | 14/242 | 186/7528 | 2.60E-03 | 3.83E-02 | 14 | HighPrior |
| hsa04380 | Osteoclast differentiation | 11/242 | 128/7528 | 2.67E-03 | 3.83E-02 | 11 | HighPrior |
| hsa04668 | TNF signaling pathway | 10/242 | 110/7528 | 2.74E-03 | 3.83E-02 | 10 | HighPrior |
| hsa04740 | Olfactory transduction | 17/68 | 448/7528 | 2.91E-07 | 3.08E-05 | 17 | LowPrior |
| hsa04742 | Taste transduction | 8/68 | 83/7528 | 6.70E-07 | 3.55E-05 | 8 | LowPrior |
| hsa04080 | Neuroactive ligand-receptor interaction | 14/68 | 338/7528 | 1.40E-06 | 4.96E-05 | 14 | LowPrior |
| hsa05320 | Autoimmune thyroid disease | 4/68 | 53/7528 | 1.28E-03 | 3.39E-02 | 4 | LowPrior |

**Figure 2.**
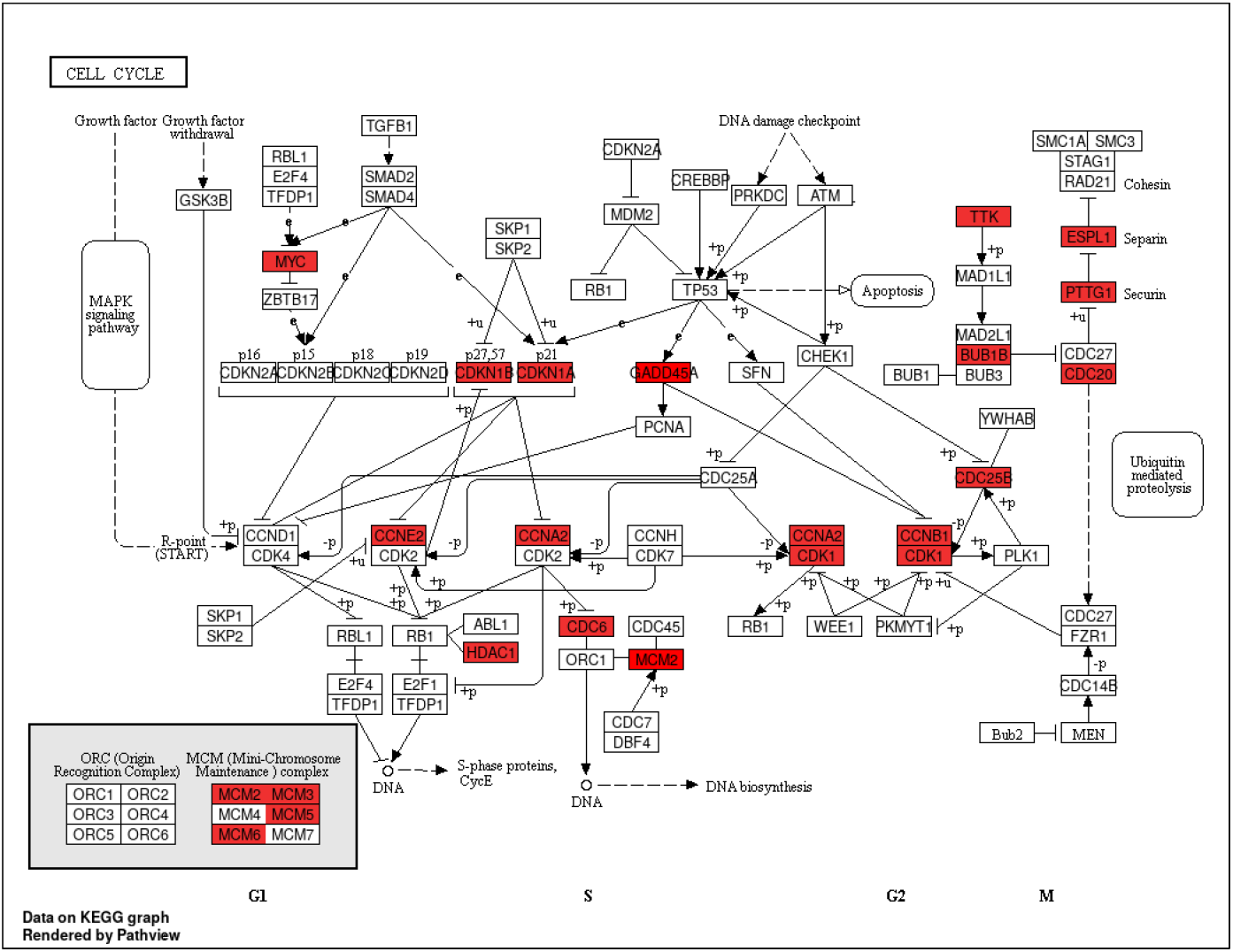
Significantly enriched Cell Cycle pathway from genes with high DE prior probabilities. The red shaded blocks indicate genes with high prior probabilities.

The 4 least frequently DE pathways include two sensory system pathways: hsa04740: Olfactory transduction and hsa04742: Taste transduction, Signaling molecules and interaction pathway. The other two significant pathways in this category were hsa04080: Neuroactive ligand-receptor interaction and hsa05320: Autoimmune thyroid disease. Our results suggest that genes belonging to these pathways show relatively stable expression across different biological conditions.

### GEOLimma method application on four validation datasets

We investigated the utility of gene-specific DE prior probabilities by performing DE analysis with GEOlimma in four evaluation datasets. Specifically, we selected two GEO series—GSE8052 and GSE15061— from platform GPL570 that enabled four DE comparisons to be made. Importantly, neither of these datasets was represented by a GEO GDS, meaning that none of the resulting comparisons were involved in DE prior probability computation. The four comparisons include Asthma vs Non-asthma (GSE8052) and three comparisons from GSE15061: Nonleukemia (Nonleuk) vs Myelodysplastic syndrome (MDS), Nonleuk vs acute myeloid leukemia (AML), and AML vs MDS. The probes for each of the datasets are represented by 20283 genes with unique mappings. Any genes without available DE prior probabilities were assigned the median value of all prior probabilities. We first identified DE genes using GEOlimma as well as the standard Limma method. This allowed us to compare the two methods, as well as characterize the extent of differential expression present in each comparison. For Limma, we considered genes to be DE if their BH-adjusted p-value ≤ 0.05. In contrast, as GEOlimma enables the calculation of a modified B score only (see Methods), we selected a B score threshold for GEOlimma significance based on the smallest Limma B score for which the Limma adjusted p-value ≤ 0.05. Using these criteria, we identified DE genes based on all relevant samples for each of the four comparisons described above. To assess the effect of small sample sizes on GEOlimma/Limma performance, we also randomly sampled 10 subsets of 40 samples (20 in each class) for each comparison and calculated the mean and standard deviation of the number of DE genes across these subsets using both methods. Table 2 lists details of each DE comparison along with summaries of our analysis results using both Limma and GEOlimma. We note that for the Asthma comparison, there are no significant DE genes based on all samples (as well as in subsets) using the Limma method. Therefore, we were not able to quantify the number of DE genes for this comparison using GEOlimma. In the remaining three comparisons, our results demonstrate that GEOlimma identifies more DE genes than Limma when applied to either all samples or 40-sample subsets. Figure 3 A helps illustrate why this is, by examining the distributions of Limma and GEOlimma B scores for the Asthma comparison. Despite the lack of significant DE genes in this comparison, use of GEOlimma results in a wider B score distribution with a marked shift to higher values compared to Limma. This difference is due to the diverse set of gene-specific DE prior probabilities used by GEOlimma, the median value of which is substantially higher than the constant value used by Limma. The potential increase in numbers of DE genes identified by GEOlimma also suggests that use of a small constant DE prior probability may result in overly conservative DE gene identification. In our PCA and t-SNE visualizations of all samples (Figures 3 B and C), we note the lack of clear separation between the Asthma and Non-asthma groups, which helps explain why no significant DE genes were detected. Figures 3 D, E, and F show the same information for a randomly selected subset of 40 samples. We note that the B scores have a similar distribution as that of all samples.

**Table 2.** Differential expression comparison details and Limma and GEOlimma DE gene counts for all samples and 10 subsets of 40 samples of each comparison.

| Datasets | Source | Sample size | Limma DEGs | GEOlimma DEGs | Limma DEGs of DE sample | GEOlimma DEGs of DE sample |
| --- | --- | --- | --- | --- | --- | --- |
| Asthma vs nonasthma | Moffatt et al. 2007 | 268 vs 136 | 0 | — | 0 | — |
| Nonleuk vs MDS | Mills et al. 2009 | 164 vs 69 | 2619 | 5823 | 98.5±161.4 | 404.3±600.9 |
| Nonleuk vs AML | Mills et al. 2009 | 202 vs 69 | 8610 | 13379 | 2788.9±901.5 | 5879.3±1415.2 |
| AML vs MDS | Mills et al. 2009 | 164 vs 202 | 10975 | 15337 | 2881.5±1068.7 | 6017±1666.7 |

**Figure 3.**
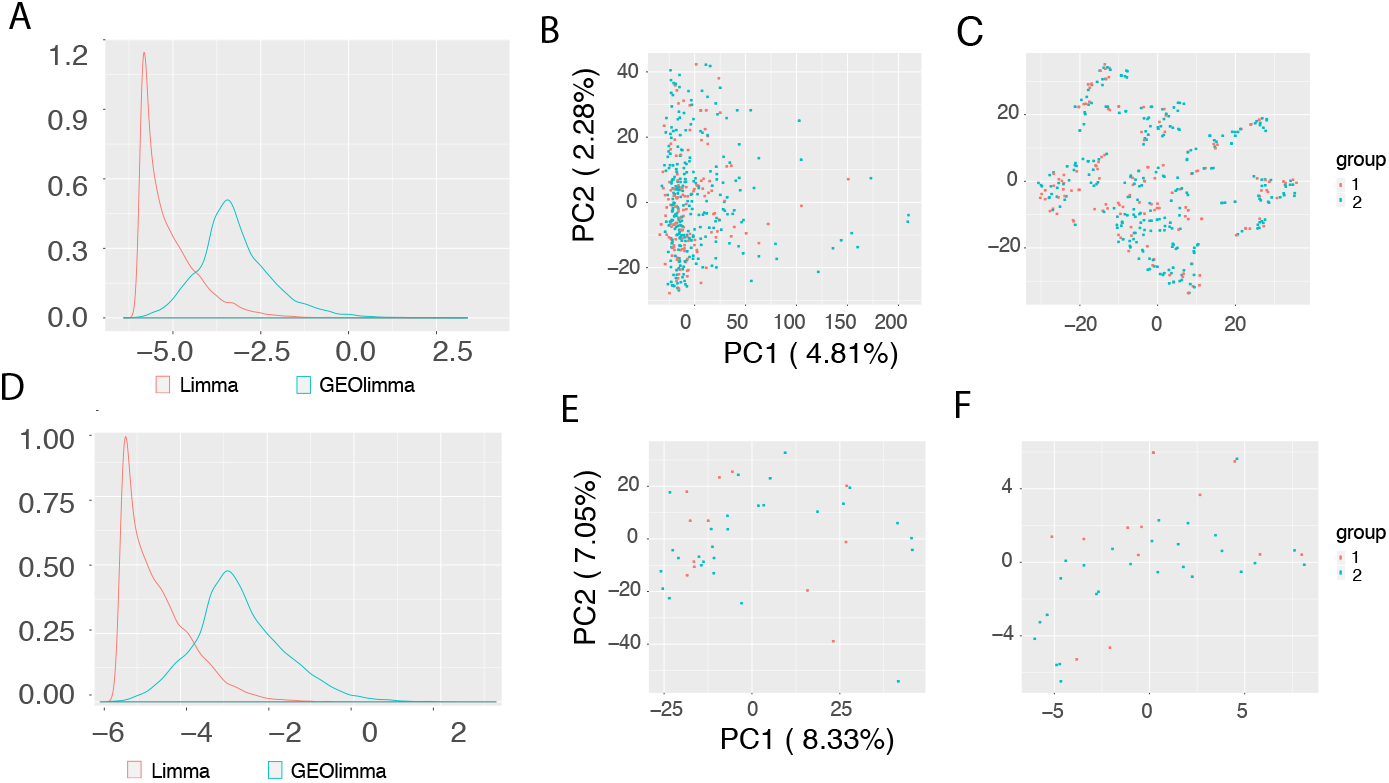
B score change and sample visualizations of asthma dataset. The top figures are generated from all samples; the bottom figures are drawn using a random subset with 40 samples. (A) and (D) depict Limma and GEOlimma B score distributions of all genes, (B) and (E) show PCA visualizations of samples, and (C) and (F) show t-SNE sample visualizations.

When looking at the top 20 most significantly DE genes for each comparison, we noted that use of GEOlimma changes the order of these genes compared to Limma, with an overall higher average B score (Figures SFigure2). To further explore this phenomenon, we counted the genes in common for the top 100 to 1000 most significantly DE genes between GEOlimma and Limma across 10 randomly selected 40-sample subsets for each comparison. The average overlap percentages were 67.3% for the Asthma comparison, 87% for Nonleuk vs MDS, while over 95% for both AML vs MDS (95.2%) and Nonleuk vs AML (95.5%)(Figure SFigure3). These results suggest that GEOlimma DE prior probabilities have a larger effect on the resulting DE gene list for datasets showing a more modest overall degree of differential expression (e.g., Asthma and Nonleuk vs MDS comparisons).

In order to explore the practical benefits of using GEOlimma, we compared the accuracy of DE gene identification between GEOlimma and Limma for each of the four DE comparisons. For each comparison, we first performed DE analysis on all samples using Limma, with the resulting significant DE genes (n = 1241 [FDR ≤0.4], 2619 [FDR ≤0.05], 8610 [FDR ≤0.05], and 10975 [FDR ≤0.05] for the Asthma, Nonleuk vs MDS, Nonleuk vs AML, and AML vs MDS comparisons) being treated as the ground truth. We note that we relaxed the significance thresholds for the Asthma comparison in order to include a sufficient number of DE genes for subsequent evaluation. Next, we randomly generated non-overlapping sample subsets for each comparison based on the minimum sample size at which the group proportions of the dataset could be maintained. For example, as GSE8052 contains 66% Asthma and 34% Non-asthma samples, the smallest sample size considered was 6 (4 Asthma, 2 Non-asthma) in order to ensure 2 samples per group. We then increased this sample size in increments of 3 to also consider subsets of 9, 12, and 15 samples. We then applied both GEOlimma and Limma on each of the sample subsets to determine which method best recovered the ground truth. Specifically, we used the R package ROCR [46] to compute areas under the receiver operating characteristic curve (AUCs) given the GEOlimma/Limma B scores and the ground truth. Figure 4 depicts the AUC improvement of GEOlimma over Limma for all four comparisons. Notably, GEOlimma consistently increases the average AUC for each of the subset sizes, with an overall average AUC improvement of 0.04. Furthermore, in the three comparisons made within GSE15061, GEOlimma increases AUC for every subset tested. Interestingly, the AUC improvement is largest for the smallest sample sizes evaluated and decreases slightly as sample size increases. This further supports the assertion that GEOlimma has a bigger impact on datasets with more modest expression differences (as would result from a small sample size). To confirm that these improvements result specifically from the DE prior probabilities learned using publicly available GPL570 data, we randomly shuffled the prior probabilities and repeated the above analysis. As seen in Figure SFfigure4, GEOlimma using randomized prior probabilities consistently decreases AUC compared to Limma.

**Figure 4.**
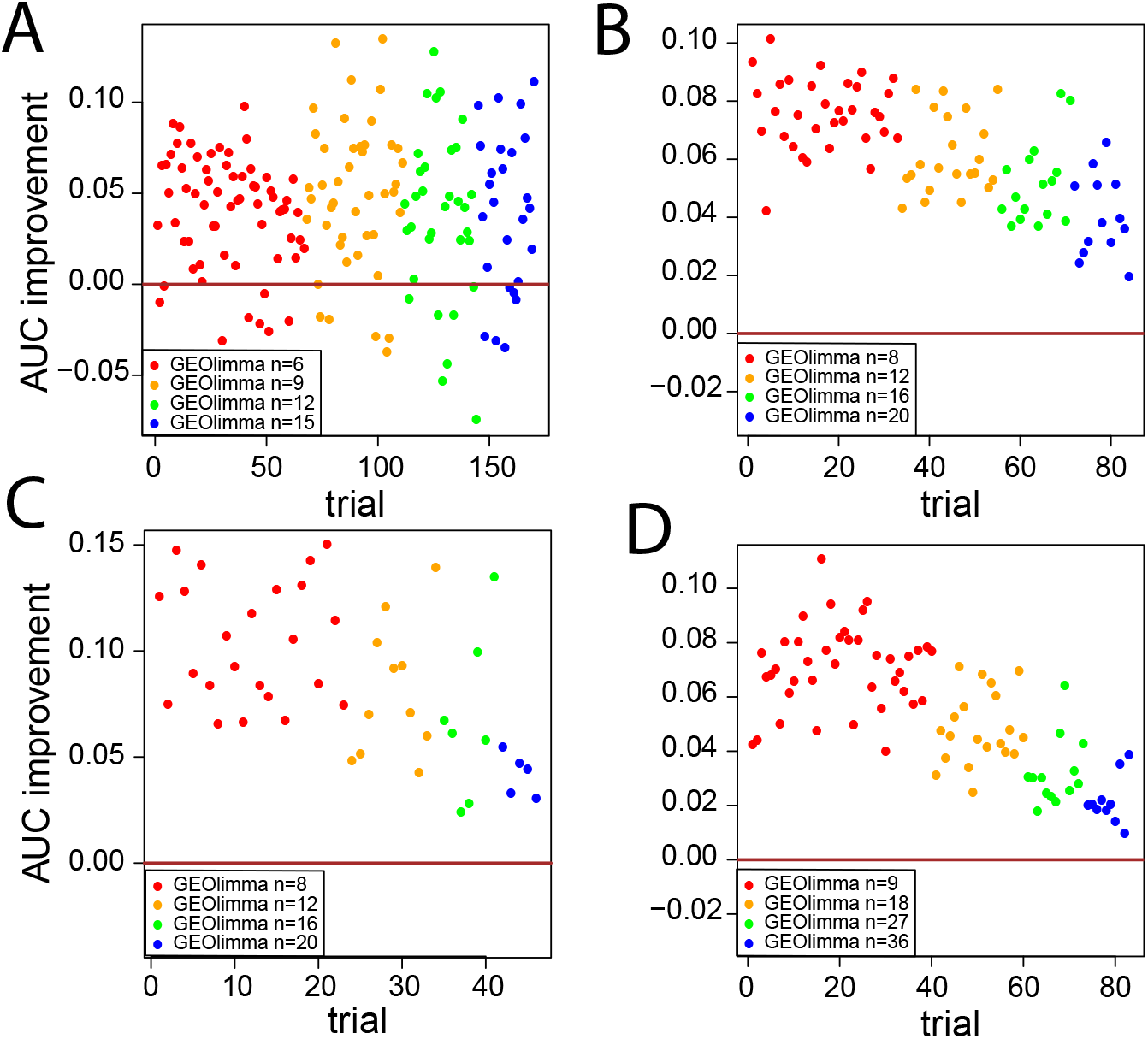
Area under the ROC curve (AUC) improvement of GEOlimma over Limma for identifying DE genes from a range of data subset sizes: A) Asthma vs Non-asthma comparison, B) Nonleukemia vs AML comparison, C) Nonleukemia vs MDS comparison, D) AML vs MDS comparison.

To quantify the experimental power gained by using GEOlimma, we converted AUC values into effective sample size. Specifically, for each of the evaluation datasets, we first calculated AUCs resulting from applying Limma to all non-overlapping sample subsets ranging in size from the minimum number needed to maintain group proportions (described above) to the total number of replicates. For example, in the Asthma comparison we considered all subsets of size 6 to 402 in increments of 3. These AUCs enabled us to fit a “standard curve” for each comparison, from which we could interpolate the mean number of samples gained by using GEOlimma given initial numbers of 6, 9, 12, and 15 (Asthma) samples. Figure SFigure5 presents the AUC standard curves and Table 3 summarizes the distribution of GEOlimma effective sample sizes for each comparison. Overall, GEOlimma leads to a substantial increase in mean effective sample sizes, particularly when applied to smaller subsets, where we observed gains of 157-288% for the smallest sample sizes evaluated for each comparison. The Asthma comparison shows the largest relative increases across all subsets, with the mean GEOlimma effective sample size more than doubling that of Limma even for the largest subset tested (m = 15). These results demonstrate the gains in experimental power for DE gene discovery that are possible with the use of GEOlimma.

**Table 3.** Distribution of effective sample sizes from applying GEOlimma to a range of data subsets (size denoted by N) for four differential expression comparisons.

| Asthma |  |  |  | AML–Nonleuk |  |  |  | Nonleuk.–MDS |  |  |  | AML–MDS |  |  |  |
| --- | --- | --- | --- | --- | --- | --- | --- | --- | --- | --- | --- | --- | --- | --- | --- |
| N | 25% | Mean | 75% | N | 25% | Mean | 75% | N | 25% | Mean | 75% | N | 25% | Mean | 75% |
| 6 | 14.39 | 23.3 | 34.83 | 8 | 19.14 | 20.55 | 22.28 | 9 | 22.1 | 24.32 | 26.53 | 10 | 22.26 | 26.44 | 30.59 |
| 9 | 21.73 | 31.98 | 45.01 | 12 | 20.98 | 22.75 | 25.03 | 18 | 27.3 | 29.54 | 31.9 | 20 | 29.67 | 34.95 | 38.25 |
| 12 | 21.72 | 29.07 | 42.2 | 16 | 23.41 | 25.64 | 26.74 | 27 | 33.35 | 35.6 | 35.72 | 30 | 38.71 | 45.13 | 48.54 |
| 15 | 16.69 | 33.57 | 49.7 | 20 | 26.03 | 27.98 | 30.17 | 36 | 41.48 | 42.53 | 42.46 | 40 | 47.95 | 50.26 | 51.5 |

### Classification performance using GEOLimma feature selection method

Feature selection is a critical step in supervised classification for diagnosis, prognosis and treatment. Here we compare the abilities of GEOlimma and Limma as feature selection methods to perform accurate classification on the four evaluation datasets. To focus on the most challenging classification tasks for each comparison, we randomly sampled subsets of size 20 from each of the two groups. Specifically, we generated 10 pairs of subsets for training, with each pair containing 40 total samples (20 per group). In the same manner, we also generated an additional 10 pairs of samples for testing. During training, we performed 10-fold cross-validation to estimate model performance. Given the large numbers of genes present in these datasets, we focused on the 1000 genes with the highest variance across all samples within each comparison. Within these 1000 genes, we selected the top 100-1000, in increments of 100, using either Limma or GEOlimma and performed classification using a logistic regression (LR) classifier. For each sampled subset, we applied a one-sided (hypothesis: GEOlimma AUC > Limma AUC) paired Wilcoxon test to compare the AUC differences between GEOlimma and Limma at each feature size (10 total). Because of the near perfect AUC observed for subsets of the AML vs MDS and Nonleuk vs AML comparisons, we only evaluated AUC differences for the Asthma and Nonleuk vs MDS comparisons using the Wilcoxon test. Table 4 shows the mean AUC differences of Asthma for each of the 10 pairs of subsets.Although many of the subsets do not show a significantly higher GEOlimma AUC, we note that the average GEOlimma - Limma AUC difference for both training and testing subsets is positive. Furthermore, subset pairs 7 and 9 show a significant GEOlimma AUC improvement in both training and testing subsets, while none of the negative AUC differences observed were significantly less than 0 (hypothesis: Limma AUC > GEOlimma AUC) in training sets. Figure 5 shows the GEOlimma and Limma AUC values at each number of features for subset pairs 7 (A) and 9 (B). For the Nonleuk vs MDS comparison, we find no significant differences between GEOlimma and Limma AUCs in training or testing subset pairs. Figure 5(C) shows one example of a training pair for this comparison. Overall, our results suggest that use of GEOlimma for feature selection can provide moderate improvements in classification performance for datasets with a modest overall degree of differential expression (e.g., Asthma comparison). For datasets with more pronounced degrees of differential expression, use of GEOlimma resulted in very similar classification performance compared to Limma.

**Table 4.** Differences in classification performance (GEOlimma AUC - Limma AUC) for 10 data subsets of the Asthma comparison. Bold p-values (Wilcoxon signed-rank test) denote statistically significant AUC improvements of GEOlimma over Limma.

| Sample order | AUCdiff | Wilcox p value | VadAUCdiff | Wilcox p value |
| --- | --- | --- | --- | --- |
| 1 | 0.0075 | 2.36E-01 | -0.01425 | 9.78E-01 |
| 2 | -0.0275 | 8.53E-01 | 0.01375 | 1.43E-01 |
| 3 | -0.0075 | 6.07E-01 | -0.025 | 9.97E-01 |
| 4 | -0.0075 | 7.79E-01 | 0.00525 | 2.39E-01 |
| 5 | -0.0175 | 9.31E-01 | 0.0105 | 3.12E-01 |
| 6 | -0.03 | 8.97E-01 | -0.0085 | 7.23E-01 |
| 7 | 0.0675 | **3.98E-02** | **0.06025** | 9.77E-04 |
| 8 | 0.025 | 1.04E-01 | -0.007 | 7.54E-01 |
| 9 | 0.0525 | **1.23E-02** | **0.0385** | 1.95E-03 |
| 10 | -0.0075 | 7.36E-01 | -0.001 | 7.93E-01 |

\*asthma dataset on LR classification

**Figure 5.**
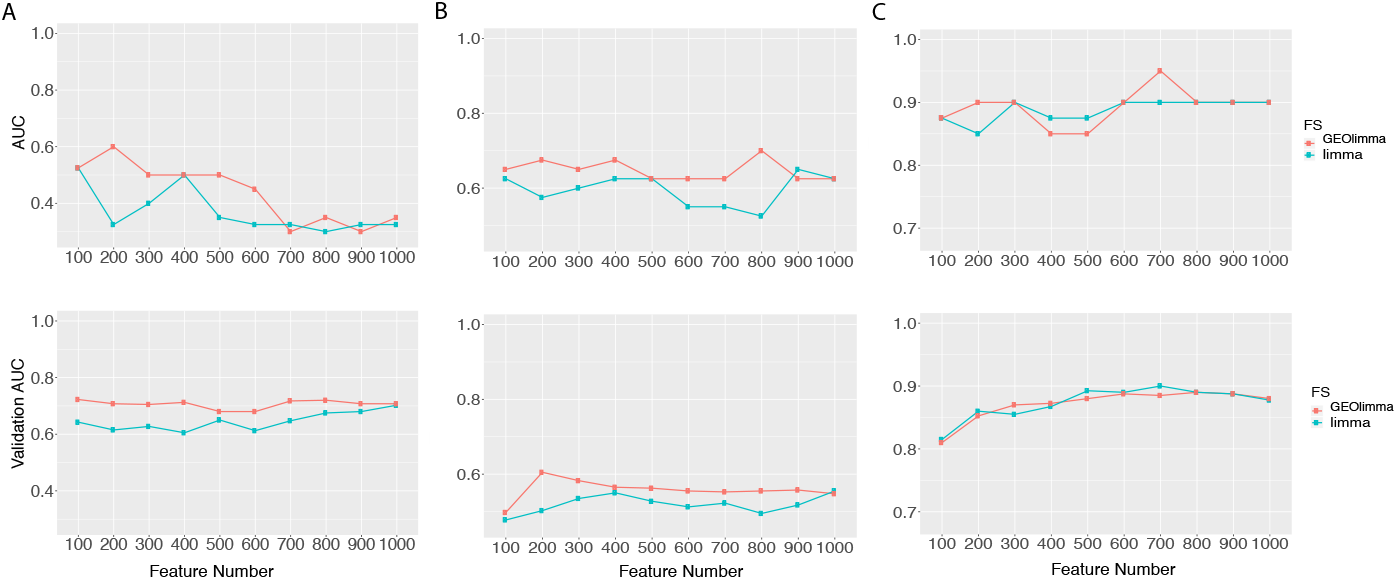
Classification performance of data subsets using a logistic regression classifier with GEOlimma and Limma feature selection methods. The x-axis indicates the number of selected features; y-axis indicates classification AUC. The top three plots display training AUC values; the bottom three plots depict validation AUCs. A) Asthma vs Non-asthma subset 7 AUCs, B) Asthma vs Non-asthma subset 9 AUCs, C) Nonleukemia vs MDS subset 9 AUCs.

## Discussion

In this study, we developed a differential expression feature selection method, GEOlimma, in which we calculated gene-level differential expression (DE) prior probabilities from large-scale GEO transcriptomics data and incorporated them into a Bayesian framework. In a DE analysis, GEOlimma detected a larger number of DE genes in four comparisons within two evaluation datasets, compared to Limma. By analyzing small sample subsets of each dataset, we showed that knowledge-driven GEOlimma substantially improved experimental power in terms of effective sample size. Furthermore, in a supervised classification analysis, GEOlimma used as a feature selection technique led to similar or better classification performance than standard Limma given noisy, small sample subsets from the Asthma comparison.

We also biologically characterized genes with especially high or low DE prior probabilities using KEGG pathway enrichment analysis. The strongest signal came from genes with high DE prior probabilities, where we detected enrichment in cell growth and death, signal transduction and cancer-related pathways. Cell growth and death are fundamental biological processes; however, deregulation of these processes is often involved in carcinosis. Specifically, resisting cell death and sustaining proliferative signaling were reported to be hallmarks of cancer [50]. This prevalence of enriched cancer-specific pathways may be indicative of an over-representation of cancer-related studies in data repositories such as GEO, which has been previously reported [51] [28]. However, while we saw excellent improvements in experimental power in differential expression analysis of three cancer-related comparisons, we note that the largest relative increases in effective sample size were observed in the Asthma comparison. This suggests that GEOlimma can also provide a substantial benefit to datasets that are unrelated to cancer.

We closely modeled GEOlimma after the widely-used differential expression analysis method Limma. Since its first publication nearly 15 years ago, papers describing the Limma method [52] [39] [38] have been cited over 10,000 times for applications in differential expression analysis of DNA microarray or RNA-Seq transcriptomics data. For the latter application, the more recently-developed voom method [52] adapts the Limma empirical Bayesian framework to read count data, which enables computation of posterior DE probabilities for RNA-Seq experiments. Although we only applied GEOlimma to DNA microarray data in this study, our approach is readily transferable to RNA-Seq data through the use of the voom methodology.

In this study, we made use of all available GPL570 GEO datasets (GDS), which we acknowledge represent a relatively small subset of all available GPL570 data at GEO. We made this selection in large part due to the high-quality curation of GDS datasets compared to the more abundant GSEs, which allowed us to easily perform multiple differential expression comparisons within each dataset. Given recent advances in natural language processing and the extraction of experimental metadata (e.g., [53]), an exciting future direction is the automatic annotation and inclusion of the larger number of GSEs (5154 for GPL570 as of June 2019) in the DE prior probability calculations. Such an expansion of a pre-existing data collection would enable subdivision and calculation of condition-specific DE prior probabilities (e.g., stem cell-related or viral infection-related), which could further improve GEOlimma performance when applied to the analysis of related datasets. One final future direction is the generalization of GEOlimma DE prior probabilities from individual values to probability distributions. In this case, DE hyperprior parameters could be calculated from pre-existing data rather than explicit prior probabilities. This modification would enable a more nuanced adjustment of DE posterior probabilities by GEOlimma given the biological characteristics of the dataset of interest.

## Conclusions

Overall, our results demonstrate that GEOlimma effectively utilized pre-existing transcriptomics data for improved differential expression and feature selection analyses. Due to its focus on gene-level differential expression, GEOlimma also has the potential to be applied to other high-throughput biological datasets.

## Declarations

### Ethics approval and consent to participate

Not applicable

## Consent for publication

Not applicable

## Availability of data and material

All datasets analyzed during the current study were previously generated and are available from Gene Expression Omnibus (GEO). Source code implementing the described analyses is available upon request.

## Authors’ contributions

All authors performed data analysis and wrote the manuscript. All authors read and approved the final manuscript.

## Acknowledgements

The authors thank the High Performance Computing Center and the Computational Research on Materials Institute at The University of Memphis (CROMIUM) for providing generous computing resources for this research.

## References

1. Harrington, C.A., Rosenow, C., Retief, J.: Monitoring gene expression using DNA microarrays. Curr. Opin. Microbiol. 3(3), 285–291 (2000)

2. Wang, Z., Gerstein, M., Snyder, M.: RNA-Seq: a revolutionary tool for transcriptomics. Nat. Rev. Genet. 10, 57 (2009)

3. Govindarajan, R., Duraiyan, J., Kaliyappan, K., Palanisamy, M.: Microarray and its applications. J. Pharm. Bioallied Sci. 4(Suppl 2), 310–2 (2012)

4. Stoughton, R.B.: Applications of DNA microarrays in biology. Annu. Rev. Biochem. 74, 53–82 (2005)

5. Van Den Berge, K., Hembach, K.M., Soneson, C., Tiberi, S., Clement, L., Love, M.I., Patro, R., Robinson, M.D.: RNA sequencing data: hitchhiker’s guide to expression analysis. Annual Review of Biomedical Data Science 2 (2018)

6. Hou, Y., Gao, B., Li, G., Su, Z.: MaxMIF: A new method for identifying cancer driver genes through effective data integration. Adv. Sci. 5(9), 1800640 (2018)

7. Alkhateeb, A., Rezaeian, I., Singireddy, S., Cavallo-Medved, D., Porter, L.A., Rueda, L.: Transcriptomics signature from Next-Generation sequencing data reveals new transcriptomic biomarkers related to prostate cancer. Cancer Inform. 18, 1176935119835522 (2019)

8. Han, J., Chen, M., Wang, Y., Gong, B., Zhuang, T., Liang, L., Qiao, H.: Identification of biomarkers based on differentially expressed genes in papillary thyroid carcinoma. Sci. Rep. 8(1), 9912 (2018)

9. Gliddon, H.D., Herberg, J.A., Levin, M., Kaforou, M.: Genome-wide host RNA signatures of infectious diseases: discovery and clinical translation. Immunology 153(2), 171–178 (2018)

10. Hira, Z.M., Gillies, D.F.: A review of feature selection and feature extraction methods applied on microarray data. Adv. Bioinformatics 2015, 198363 (2015)

11. Nazarov, P.V., Muller, A., Kaoma, T., Nicot, N., Maximo, C., Birembaut, P., Tran, N.L., Dittmar, G., Vallar, L.: RNA sequencing and transcriptome arrays analyses show opposing results for alternative splicing in patient derived samples. BMC Genomics 18(1), 443 (2017)

12. Wang, Y., Barbacioru, C., Hyland, F., Xiao, W., Hunkapiller, K.L., Blake, J., Chan, F., Gonzalez, C., Zhang, L., Samaha, R.R.: Large scale real-time PCR validation on gene expression measurements from two commercial long-oligonucleotide microarrays. BMC Genomics 7, 59 (2006)

13. Chen, J.J., Hsueh, H.-M., Delongchamp, R.R., Lin, C.-J., Tsai, C.-A.: Reproducibility of microarray data: a further analysis of microarray quality control (MAQC) data. BMC Bioinformatics 8, 412 (2007)

14. Wei, C., Li, J., Bumgarner, R.E.: Sample size for detecting differentially expressed genes in microarray experiments. BMC Genomics 5, 87 (2004)

15. Clarke, R., Ressom, H.W., Wang, A., Xuan, J., Liu, M.C., Gehan, E.A., Wang, Y.: The properties of high-dimensional data spaces: implications for exploring gene and protein expression data. Nat. Rev. Cancer 8(1), 37–49 (2008)

16. Boluki, S., Esfahani, M.S., Qian, X., Dougherty, E.R.: Incorporating biological prior knowledge for bayesian learning via maximal knowledge-driven information priors. BMC Bioinformatics 18(Suppl 14), 552 (2017)

17. McNeish, D.: On using bayesian methods to address small sample problems. Struct. Equ. Modeling 23(5), 750–773 (2016)

18. Ashburner, M., Ball, C.A., Blake, J.A., Botstein, D., Butler, H., Cherry, J.M., Davis, A.P., Dolinski, K., Dwight, S.S., Eppig, J.T., Harris, M.A., Hill, D.P., Issel-Tarver, L., Kasarskis, A., Lewis, S., Matese, J.C., Richardson, J.E., Ringwald, M., Rubin, G.M., Sherlock, G.: Gene ontology: tool for the unification of biology. the gene ontology consortium. Nat. Genet. 25(1), 25–29 (2000)

19. The Gene Ontology Consortium: The gene ontology resource: 20 years and still GOing strong. Nucleic Acids Res. 47(D1), 330–338 (2019)

20. Kanehisa, M., Goto, S.: KEGG: kyoto encyclopedia of genes and genomes. Nucleic Acids Res. 28(1), 27–30 (2000)

21. Kanehisa, M., Furumichi, M., Tanabe, M., Sato, Y., Morishima, K.: KEGG: new perspectives on genomes, pathways, diseases and drugs. Nucleic Acids Res. 45(D1), 353–361 (2017)

22. Daigle, B.J. Jr, Altman, R.B.: M-BISON: microarray-based integration of data sources using networks. BMC Bioinformatics 9, 214 (2008)

23. Morrison, J.L., Breitling, R., Higham, D.J., Gilbert, D.R.: GeneRank: using search engine technology for the analysis of microarray experiments. BMC Bioinformatics 6, 233 (2005)

24. Edgar, R., Domrachev, M., Lash, A.E.: Gene expression omnibus: NCBI gene expression and hybridization array data repository. Nucleic Acids Res. 30(1), 207–210 (2002)

25. Barrett, T., Wilhite, S.E., Ledoux, P., Evangelista, C., Kim, I.F., Tomashevsky, M., Marshall, K.A., Phillippy, K.H., Sherman, P.M., Holko, M., Yefanov, A., Lee, H., Zhang, N., Robertson, C.L., Serova, N., Davis, S., Soboleva, A.: NCBI GEO: archive for functional genomics data sets–update. Nucleic Acids Res. 41(Database issue), 991–5 (2013)

26. Kolesnikov, N., Hastings, E., Keays, M., Melnichuk, O., Tang, Y.A., Williams, E., Dylag, M., Kurbatova, N., Brandizi, M., Burdett, T., Megy, K., Pilicheva, E., Rustici, G., Tikhonov, A., Parkinson, H., Petryszak, R., Sarkans, U., Brazma, A.: ArrayExpress update–simplifying data submissions. Nucleic Acids Res. 43(Database issue), 1113–6 (2015)

27. Daigle, B.J. Jr, Deng, A., McLaughlin, T., Cushman, S.W., Cam, M.C., Reaven, G., Tsao, P.S., Altman, R.B.: Using pre-existing microarray datasets to increase experimental power: application to insulin resistance. PLoS Comput. Biol. 6(3), 1000718 (2010)

28. Engreitz, J.M., Daigle, B.J. Jr, Marshall, J.J., Altman, R.B.: Independent component analysis: mining microarray data for fundamental human gene expression modules. J. Biomed. Inform. 43(6), 932–944 (2010)

29. Kim, R.D., Park, P.J.: Improving identification of differentially expressed genes in microarray studies using information from public databases. Genome Biol. 5(9), 70 (2004)

30. Chen, R., Morgan, A.A., Dudley, J., Deshpande, T., Li, L., Kodama, K., Chiang, A.P., Butte, A.J.: FitSNPs: highly differentially expressed genes are more likely to have variants associated with disease. Genome Biol. 9(12), 170 (2008)

31. Crow, M., Lim, N., Ballouz, S., Pavlidis, P., Gillis, J.: Predictability of human differential gene expression. Proc. Natl. Acad. Sci. U. S. A. 116(13), 6491–6500 (2019)

32. Saeys, Y., Inza, I., Larrañaga, P.: A review of feature selection techniques in bioinformatics. Bioinformatics 23(19), 2507–2517 (2007)

33. He, Z., Yu, W.: Stable feature selection for biomarker discovery. Comput. Biol. Chem. 34(4), 215–225 (2010)

34. Bolón-Canedo, V., Sánchez-Maroño, N., Alonso-Betanzos, A., Benítez, J.M., Herrera, F.: A review of microarray datasets and applied feature selection methods. Inf. Sci. 282, 111–135 (2014)

35. Ang, J.C., Mirzal, A., Haron, H., Hamed, H.N.A.: Supervised, unsupervised, and Semi-Supervised feature selection: A review on gene selection. IEEE/ACM Trans. Comput. Biol. Bioinform. 13(5), 971–989 (2016)

36. Abusamra, H.: A comparative study of feature selection and classification methods for gene expression data of glioma. Procedia Comput. Sci. 23, 5–14 (2013)

37. Smyth, G.K.: Linear models and empirical bayes methods for assessing differential expression in microarray experiments. Stat. Appl. Genet. Mol. Biol. 3, 3 (2004)

38. Smyth, G.K.: limma: Linear models for microarray data. In: Gentleman, R., Carey, V.J., Huber, W., Irizarry, R.A., Dudoit, S. (eds.) Bioinformatics and Computational Biology Solutions Using R and Bioconductor, pp. 397–420. Springer, New York, NY (2005)

39. Ritchie, M.E., Phipson, B., Wu, D., Hu, Y., Law, C.W., Shi, W., Smyth, G.K.: limma powers differential expression analyses for RNA-sequencing and microarray studies. Nucleic Acids Res. 43(7), 47 (2015)

40. Benjamini, Y., Hochberg, Y.: Controlling the False Discovery Rate: A Practical and Powerful Approach to Multiple Testing (1995)

41. Moffatt, M.F., Kabesch, M., Liang, L., Dixon, A.L., Strachan, D., Heath, S., Depner, M., von Berg, A., Bufe, A., Rietschel, E., Heinzmann, A., Simma, B., Frischer, T., Willis-Owen, S.A.G., Wong, K.C.C., Illig, T., Vogelberg, C., Weiland, S.K., von Mutius, E., Abecasis, G.R., Farrall, M., Gut, I.G., Lathrop, G.M., Cookson, W.O.C.: Genetic variants regulating ORMDL3 expression contribute to the risk of childhood asthma. Nature 448(7152), 470–473 (2007)

42. Mills, K.I., Kohlmann, A., Williams, P.M., Wieczorek, L., Liu, W.-M., Li, R., Wei, W., Bowen, D.T., Loeffler, H., Hernandez, J.M., Hofmann, W.-K., Haferlach, T.: Microarray-based classifiers and prognosis models identify subgroups with distinct clinical outcomes and high risk of AML transformation of myelodysplastic syndrome. Blood 114(5), 1063–1072 (2009)

43. Yu, G., Wang, L.-G., Han, Y., He, Q.-Y.: clusterprofiler: an R package for comparing biological themes among gene clusters. OMICS 16(5), 284–287 (2012)

44. Luo, W., Brouwer, C.: Pathview: an R/Bioconductor package for pathway-based data integration and visualization. Bioinformatics 29(14), 1830–1831 (2013)

45. Maaten, L.v.d., Hinton, G.: Visualizing data using t-SNE. J. Mach. Learn. Res. 9(Nov), 2579–2605 (2008)

46. Sing, T., Sander, O., Beerenwinkel, N., Lengauer, T.: ROCR: visualizing classifier performance in R. Bioinformatics 21(20), 3940–3941 (2005)

47. Pedregosa, F., Varoquaux, G., Gramfort, A., Michel, V., Thirion, B., Grisel, O., Blondel, M., Prettenhofer, P., Weiss, R., Dubourg, V., Vanderplas, J., Passos, A., Cournapeau, D., Brucher, M., Perrot, M., Duchesnay, É.: Scikit-learn: Machine learning in python. J. Mach. Learn. Res. 12(Oct), 2825–2830 (2011)

48. Trevor, H., Robert, T., Jh, F.: The elements of statistical learning: data mining, inference, and prediction. New York, NY: Springer (2009)

49. Pandurangan, A.K., Divya, T., Kumar, K., Dineshbabu, V., Velavan, B., Sudhandiran, G.: Colorectal carcinogenesis: Insights into the cell death and signal transduction pathways: A review. World J. Gastrointest. Oncol. 10(9), 244–259 (2018)

50. Hanahan, D., Weinberg, R.A.: Hallmarks of cancer: the next generation. Cell 144(5), 646–674 (2011)

51. Huttenhower, C., Haley, E.M., Hibbs, M.A., Dumeaux, V., Barrett, D.R., Coller, H.A., Troyanskaya, O.G.: Exploring the human genome with functional maps. Genome Res. 19(6), 1093–1106 (2009)

52. Law, C.W., Chen, Y., Shi, W., Smyth, G.K.: voom: Precision weights unlock linear model analysis tools for RNA-seq read counts. Genome Biol. 15(2), 29 (2014)

53. Giles, C.B., Brown, C.A., Ripperger, M., Dennis, Z., Roopnarinesingh, X., Porter, H., Perz, A., Wren, J.D.: ALE: automated label extraction from GEO metadata. BMC Bioinformatics 18(Suppl 14), 509 (2017)

